# E93-depleted adult insects preserve the prothoracic gland and molt again

**DOI:** 10.1101/2020.03.01.971580

**Authors:** Orathai Kamsoi, Xavier Belles

## Abstract

Insect metamorphosis originated around the middle Devonian, associated with the innovation of the final molt; this occurs after the histolysis of the prothoracic gland (PG; which produces the molting hormone) in the first days of adulthood. We previously hypothesized that transcription factor E93 was crucial in the emergence of metamorphosis, since it triggers metamorphosis in extant insects. This work on the cockroach *Blattella germanica* reveals that E93 also plays a crucial role in the histolysis of PG, which fits the above hypothesis. Previous studies have shown that the transcription factor FTZ-F1 is essential for PG histolysis. We have found that FTZ-F1 depletion, towards the end of the final nymphal instar, downregulates the expression of *E93*, while E93-depleted nymphs molt to adults that retain a functional PG. Interestingly, these adults are able to molt again, which is exceptional in insects. The study of insects able to molt again in the adult stage may reveal clues as to how nymphal epidermal cells definitively become adult cells, and if it is possible to revert this process.

**Summary statement:** The prothoracic gland disintegrates after insect metamorphosis. It was believed that the factor FTZ-F1 determines this disintegration. This work reveals that FTZ-F1 action is mediated by the factor E93.

## INTRODUCTION

One of the most successful innovations in insect evolution is metamorphosis, as shown by the fact that more than 95% of extant insects develop through this type of postembryonic transformation. The simplest mode of metamorphosis, hemimetaboly, or incomplete metamorphosis, originated with the clade Pterygota (viz. the presence of wings), around the middle Devonian, ca. 400 Mya. Subsequently, during the early Carboniferous, ca. 350 Mya, holometaboly, or complete metamorphosis, evolved from hemimetaboly (Belles, 2019b; Belles, 2020). Consubstantial with the emergence of wings and hemimetabolan metamorphosis is the innovation of the final molt, mainly achieved through the histolysis of the prothoracic gland (PG; the gland that produces the molting hormone), which takes place after the winged and reproductively competent adult stage has been reached.

The molecular mechanism that regulates insect metamorphosis is condensed in the MEKRE93 pathway (Belles and Santos, 2014), through which juvenile hormone (JH) bound to its receptor Methoprene tolerant (Met), and induces the expression of Krüppel homolog 1 (Kr-h1); this, in turn, represses the expression of E93. The players most directly involved in regulating metamorphosis are Kr-h1, the transducer of the antimetamorphic signal of JH, and E93, the master trigger of metamorphosis (see (Belles, 2019a)). *E93* was originally discovered as an ecdysone-induced late prepupal specific gene, during research into the histolysis of the salivary glands in *Drosophila melanogaster* metamorphosis, a process in which E93 plays a key role (Baehrecke and Thummel, 1995; Lee et al., 2000; Woodard et al., 1994). Subsequently, Mou et al. (2012) found that *E93* is widely expressed in adult cells that form in the pupa of *D. melanogaster*, being required for morphogenesis patterning processes. Further experiments revealed that E93-depleted larvae of *D. melanogaster* are able to pupate but die at the end of the pupal stage. Similar results were observed in the beetle *Tribolium castaneum*, where E93 depletion prevented the pupal-adult transition, resulting in the formation of a supernumerary second pupa (Ureña et al., 2014). Moreover, studies on the cockroach *Blattella germanica* have revealed that E93 depletion prevents the nymph-adult transition, resulting in reiterated supernumerary nymphal instars (Ureña et al., 2014), and that the expression of *E93* in juvenile nymphs is inhibited by Kr-h1, which establishes the MEKRE93 pathway (Belles and Santos, 2014).

Importantly, the MEKRE93 pathway, including the role of E93 as a metamorphosis trigger, is conserved in extant metamorphosing insects (Belles, 2020). This suggests that it was operative in the pterygote last common ancestor. If this is true, then a single mechanism, the MEKRE93 pathway and E93 in particular, might have simultaneously promoted metamorphosis, including wing maturation, and PG histolysis (and hence the final molt), at the origin of the Pterygota (Belles, 2019b). The role of E93 as a promoter of metamorphosis and wing maturation has been thoroughly demonstrated (Belles and Santos, 2014; Mou et al., 2012; Ureña et al., 2014; Uyehara et al., 2017). In contrast, nothing is known about the possible role of E93 in PG histolysis. The aim of this work is to explore this possibility using *B. germanica* as a model.

Histolysis of cells and tissues through programmed cell death (PCD) is consubstantial with metamorphosis, especially in holometabolans, as during this process, new structures are generated, while others disappear. In this way, some cells and tissues disintegrate (like the salivary glands), whereas others undergo remodeling, with partial cell replacement (like in the fat body) (Tettamanti and Casartelli, 2019). Intensive studies based on the larva-pupa transition of *D. melanogaster*, have demonstrated the determinant role of ecdysone signaling in PCD processes that affect the salivary glands, midgut and fat body. Ecdysone, or rather its most bioactive derivative, 20-hydroxyecdysone (20E), binds the receptor, EcR, forms a complex with the coreceptor retinoid X receptor (RXR), and initiates a gene expression cascade (Hill et al., 2013) that includes *fushi tarazu-factor 1* (*ftz-f1*) and *E93* among the most important genes in the context of PCD (Baehrecke and Thummel, 1995; Broadus et al., 1999). Next, the corresponding proteins promote the expression of PCD genes, like *reaper* (*rpr*) and *head involution defective* (*hid*), as well as caspases and other direct PCD mediators (Jiang et al., 2000; Lee et al., 2000). Other important players are the inhibitors of apoptosis (IAPs), which protect cells and tissues from PCD by preventing caspase activity (Orme and Meier, 2009), although their inhibitory activity on PCD is counteracted by Rpr and Hid (Martin, 2002). Therefore, at the larva-pupa transition, downregulation of *iap1* expression provides the competence for PCD triggered by cell death factors (Orme and Meier, 2009; Yin and Thummel, 2005; Yin et al., 2007).

Although E93 had never been related to the PG disintegration, it has been characterized as promoting PCD during metamorphosis in various tissues and species. The most thorough studies have been carried out on the salivary glands of *D. melanogaster*, where stage- and tissue-specific expression of E93 triggers their histolysis (Baehrecke and Thummel, 1995; Berry and Baehrecke, 2007; Lee et al., 2000; Lee et al., 2002b; Woodard et al., 1994). The action of E93 on PCD during *D. melanogaster* metamorphosis has also been reported to occur in the midgut (Lee and Baehrecke, 2001; Lee et al., 2002a) and fat body (Liu et al., 2014). More recently, E93 has been shown to play the same PCD role in the fat body of the silkworm *Bombyx mori* (Liu et al., 2015).

In *B. germanica*, our model, previous studies have revealed that *iap1* expression in the PG declines during the nymph-adult transition, which is followed by PG histolysis. In *B. germanica*, PG histolysis is regulated by 20E signaling, which leads to a dramatic upregulation of *ftz-f1* expression, the gene product of this plays a crucial role in the PCD process (Mané-Padrós et al., 2010). Given the above antecedents, in this work we conjectured that the PCD action of FTZ-F1 on the PG might be mediated by E93. This presumption has been confirmed, as FTZ-F1 depletion towards the end of the final nymphal instar downregulates *E93* expression, while E93-depletion results in adults that retain a functional PG. It was still more intriguing, however, to observe that the E93-depleted adults were able to molt again. Studying this phenotype could provide important clues as to how nymphal epidermal cells definitively become adult cells, and if it is possible to revert that process.

## RESULTS

### *E93* expression is tissue-specific

We studied the expression pattern of E93 in various tissues involved in metamorphosis, namely the PG, the corpora cardiaca-corpora allata (CC-CA) complex, the epidermal tissue (represented by the abdominal tergites two to seven), and the wing pads. We studied the three most important premetamorphic instars, the fourth (antepenultimate) nymphal instar (N4), the penultimate nymphal instar (N5) and the final nymphal instar (N6). At the same time, we measured *ftz-f1* expression given its relationship with PG histolysis. The PG patterns are clearly differentiated from those of other tissues (Fig. 1). The highest expression levels of *E93* in the CC-CA complex, epidermis, and wing pads are found towards the middle of N6, while in the PG they peak in N6D8. As for *ftz-f1*,there is a single expression peak in N6D8 in the PG (coinciding with that of *E93*), while in the other tissues the expression levels fluctuate as a result of the pulses of 20E that occur around each molt (Fig. 1).

**Fig. 1.**
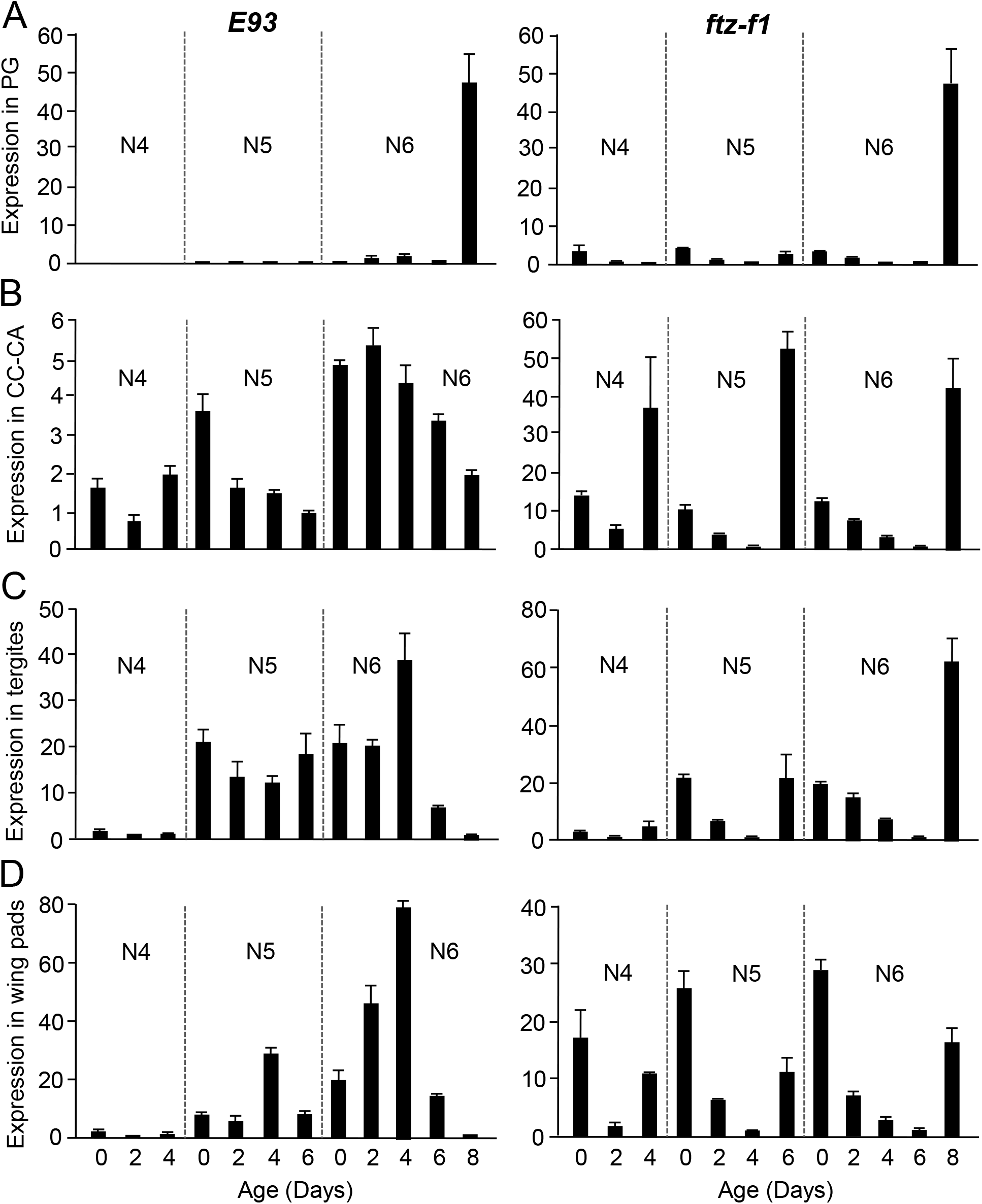
*E93* and *ftz-f1* expression in the last nymphal instars of *Blattella germanica*. (A) Expression in the prothoracic gland (PG). (B) Expression in corpora cardiaca-corpora allata (CC-CA) complex. (C) Expression in tergites. (D) Expression in wing pads. The expression was measured during the fourth (antepenultimate) nymphal instar (N4), the fifth (penultimate) nymphal instar (N5) and the sixth (final) nymphal instar (N6) in female insects. The results are indicated as copies of the examined mRNA per 1000 copies of BgActin-5c mRNA, and are expressed as the mean ± SEM (n=3).

### *E93* interference late in the final nymphal instar triggers the formation of adults that molt once again

To maximize the depletion of *E93* transcripts in the PG while only minimally affecting the other metamorphic tissues, we injected dsRNA targeting *E93* (dsE93) into N6D6 insects. At this stage, the of *E93* expression peak had already occurred in the CC-CA complex, tergites, and wing pads, while it was still to come in the PG. Insects injected in N6D6 with either dsMock (control) (n = 67) or dsE93 (n = 132) all molted to adults two days later. The insects from the control experiments presented a normal adult external morphology, with well-shaped and completely extended forewings and hindwings (Fig. 2A). As for the dsE93-treated nymphs, 85.6% molted to adults presenting a normal external appearance, with well-shaped and completely extended forewings and hindwings, and 14.4% molted to adults with the wings only partially extended, particularly the hindwings (Fig. 2A).

**Fig. 2.**
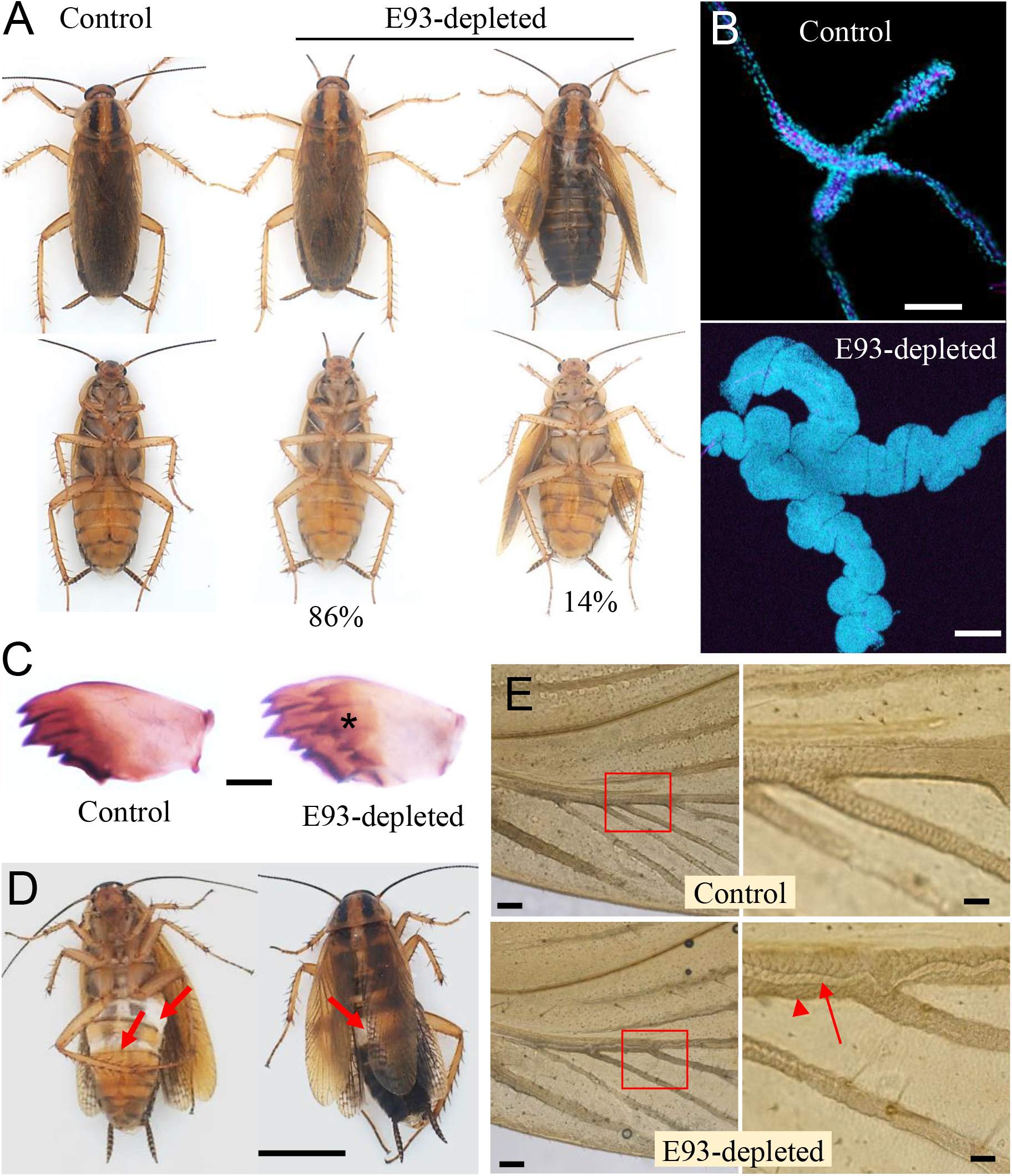
Effect of *E93* mRNA depletion in the imaginal molt of *Blattella germanica*. (A) Dorsal and ventral views of control and E93-depleted adults. Sixth instar females nymphs were treated on day 6 with dsMock (control) or dsE93, and the morphology after the imaginal molt was examined; the percentages of E93-depleted adults with extended and unextended wings are indicated; scale bar: 5 mm. (B) Prothoracic gland of control and E93-depleted adults on day 8 of adult life; the glands were stained with phalloidin (pink) and DAPI (blue); scale bar: 0.1 mm. (C) Right mandible of control and E93-depleted adults on day 9 of adult life; note the new mandible formed under the old one in the E93-depleted adults after apolysis (asterisk); scale bar: 0.2 mm. (D) E93-depleted adults attempting to molt on day 10 of adult life; the arrows show areas where the old cuticle separated, showing the new cuticle formed after the apolysis; scale bar: 5 mm. (E) Basal part of the left forewing of control and E93-depleted adults on day 9 of adult life; the part indicated with the red square is shown at higher magnification in the panels on the right; note the new veins (arrow) formed under the old ones (arrowhead) in the E93-depleted adults after apolysis; scale bar: 0.2 mm (left panels), 0.05 mm (right panels).

It has been shown previously that the adult of *B. germanica* start disintegrating the PG after the imaginal molt, and three days later the disintegration is very apparent, since only the axes of muscle, tracheae and nerve of the X-shaped gland, and a few secretory cells, are observed (Romaña et al., 1995). In our RNAi experiments, control adults culminated the histolysis and disintegration process in 3 to 5 days, while the E93-depleted adults retained the integrity of the PG, as observed 8 days after the imaginal molt, when its general morphology resembles that of an N6 PG (Fig. 2B). There is an intriguing difference in cell proliferation: normally, N6 PGs undergo cell division during the first two days of the instar (Kamsoi and Belles, 2019), whereas no cell division was observed in the PG of E93-depleted adults (Fig. S1).

After the imaginal molt, the E93-depleted adults started to molt again, the first symptoms of which were detected 8 days after that molt, when shadows of the new cuticular structures, including new mandibles, could be observed by transparency under the old cuticle. On day 9 the new mandibles were clearly defined (Fig. 2C). One day later, the E93-depleted adults tried to undertake ecdysis, which could not be completed as they were unable to shed the exuvia (Fig. 2D). Even the wings undertook a new apolysis, and a new set of wing veins could be seen by transparency under the veins of the E93-depleted adults, being most clearly visible in the forewings (Fig. 2E).

### Molting of E93-depleted adults results from the effective depletion of E93 in the PG, which remains functional

We determined the efficiency of the RNAi treatment by measuring the E93 transcript decrease 48 h after the dsRNA treatment, i.e., in N6D8. Results showed that E93 mRNA levels were significantly depleted (ca. 77%) in the PG (Fig. 3A). E93 transcript depletion was also observed in the other tissues studied (CC-CA, tergites, and wing pads), although the baseline levels were low, as expected from the expression patterns (Fig. 3A). Simultaneous measurement of *ftz-f1* mRNA levels indicated that E93 depletion did not affect *ftz-f1* expression in the PG, but did trigger a dramatic up-regulation in the CC-CA, tergites and wing pads (Fig. 3A). We also observed that the interference of E93 in the late final nymphal instar did not significantly affect the expression in epidermal tissues of *Kr-h1* and *BR-C*, two important genes in adult morphogenesis (Fig. 3B).

**Fig. 3.**
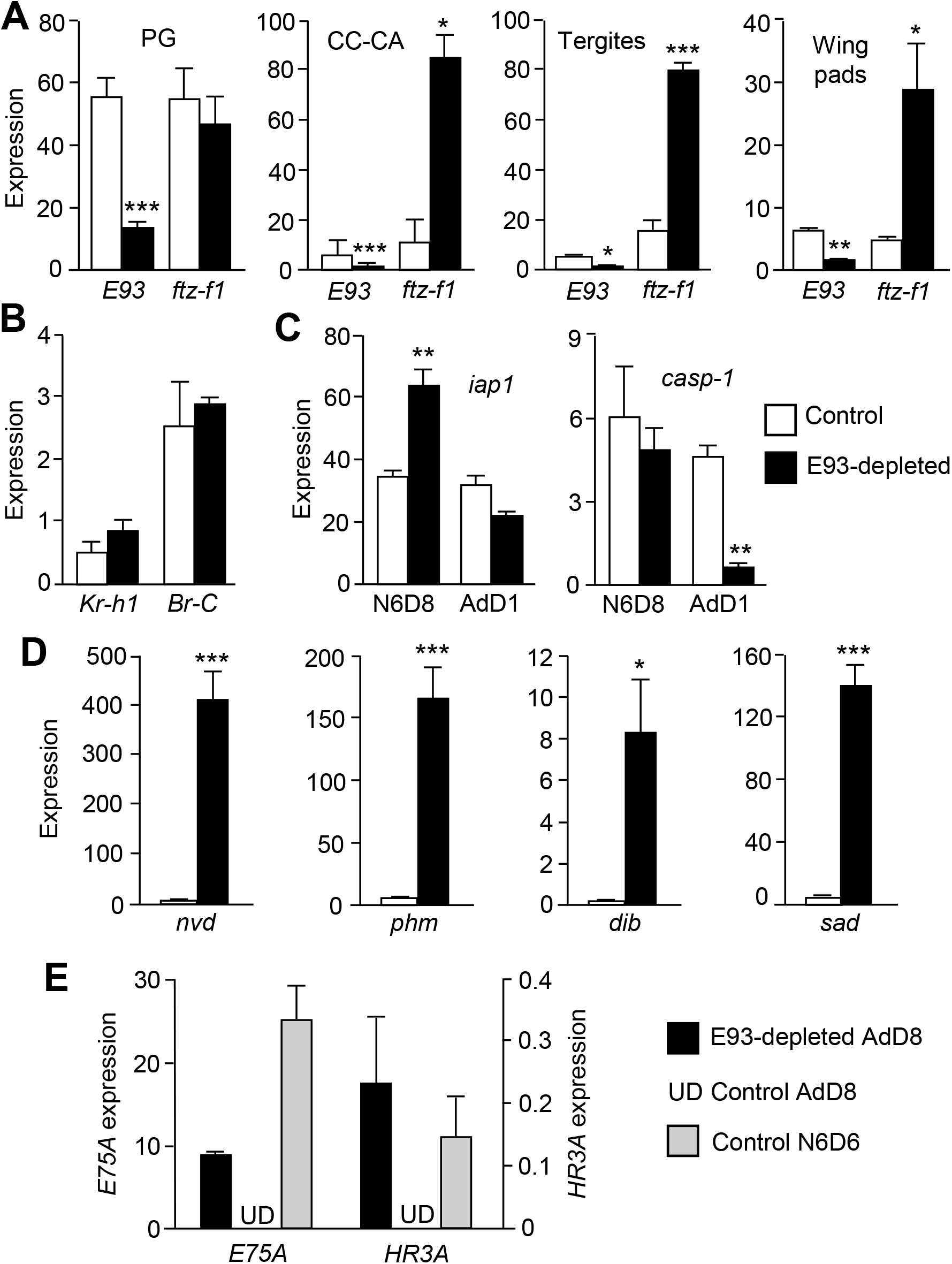
Effect of E93 depletion on gene expression in *Blattella germanica*. (A) Expression of *E93* and *ftz-f1* in the prothoracic gland (PG), corpora cardiaca-corpora allata (CC-CA) complex, tergites, and wing pads from E93-depleted insects and controls on day 8 of the sixth nymphal instar (N6D8). (B) Expression of *Kr-h1* and *BR-C* in tergites from E93-depleted insects and controls in N6D8. (C) Expression of *iap1* and *casp-1* in the PG of E93-depleted adults and controls in N6D8 and on the first day of the adult stage (AdD1). (D) Expression of the steroidogenic genes *nvd, phm, dib*, and *sad*, in the PG from E93-depleted adults and controls in AdD8. (E) Expression of the ecdysteroid signaling genes *E75A* and *HR3A* in the PG from E93-depleted adults and controls in AdD8, and in control nymphs in N6D6. The results are indicated as copies of the examined mRNA per 1000 copies of BgActin-5c mRNA, and expressed as the mean ± SEM (n=3); the asterisks indicate statistically significant differences with respect to the controls (*p<0.05, **p<0.01, ***p<0.001) according to the student’s *t*-test. In panel E “UD”means that mRNA levels were under the detection limit.

We measured the expression of two genes with opposite functions in PG histolysis, *iap1* and *caspase-1* (*casp-1*), on the last day of the final nymphal instar (N6D8), and the first day of the adult stage (AdD1). The results showed that in N6D8 *iap1* expression is significantly higher in E93-depleted insects, although in AdD1 this returns to levels similar to those seen in the controls. In contrast, *casp-1* expression in AdD1 is significantly lower than that found in the controls (Fig. 3C).

In addition, we measured the expression of the steroidogenic genes *neverland* (*nvd*), *phantom* (*phm*), *disembodied* (*dib*) and *shadow* (*sad*) in the PG of control and E93-depleted 8-day-old adults (AdD8). All the genes were found to be efficiently expressed in the PG of E93-depleted adults, whereas, as expected, expression was very low or undetectable in the controls (Fig. 3D). Consistently, the ecdysone-dependent genes *E75A* and *HR3A* were significantly expressed in the PG of E93-depleted AdD8, suggesting that the gland was producing and was exposed to ecdysteroids, whereas the expression of these two genes was undetectable in the PG of the AdD8 controls (Fig. 3E). Indeed, the expression of *E75A* and *HR3A* in the PG of E93-depleted AdD8 is relatively comparable to that measured in untreated N6D6 (Fig. 3E), the age of the final nymphal instar at which the ecdysone pulse is produced (Cruz et al., 2003).

### PG death induced by FTZ-F1 is mediated by E93

We had previously reported that FTZ-F1 plays a critical role in the histolysis of PG in *B. germanica* (Mané-Padrós et al., 2010). Given that E93 depletion in the PG prevented gland histolysis (Fig. 2C), we wondered about the relationships between FTZ-F1 and E93 in the process. As discussed above, E93 appears to repress *ftz-f1* expression in CC-CA, tergites and wing pads, but not in the PG (Fig. 3A). A possible explanation for our observations on the action of E93 and FTZ-F1 on PG histolysis is to consider that FTZ-F1 would enhance the expression of E93 in the PG, and that E93 would be the gland histolysis effector. To test this conjecture, we treated N6D7 female nymphs (i.e., one day before the peak of *ftz-f1* and *E93* in the PG, Fig. 1) with dsFTZ-F1. We then measured the expression of *E93* (and *ftz-f1*) in the PG. The results showed that the RNAi of *ftz-f1* was efficient, as 6 h after the treatment the corresponding mRNA levels became significantly reduced in the PG (as well as in the CC-CA). At 12 h, *ftz-f1* mRNA levels were still decreased in the PG, but the differences with respect to the controls were not statistically significant. Importantly, the mRNA levels of *E93* became significantly down-regulated in the PG at 6 and 12 h, but not in the CC-CA. At 24 h, *ftz-f1* mRNA levels in the PG recovered normal (control) levels, but the significant downregulation of *E93* expression persisted (Fig. 4).

**Fig. 4.**
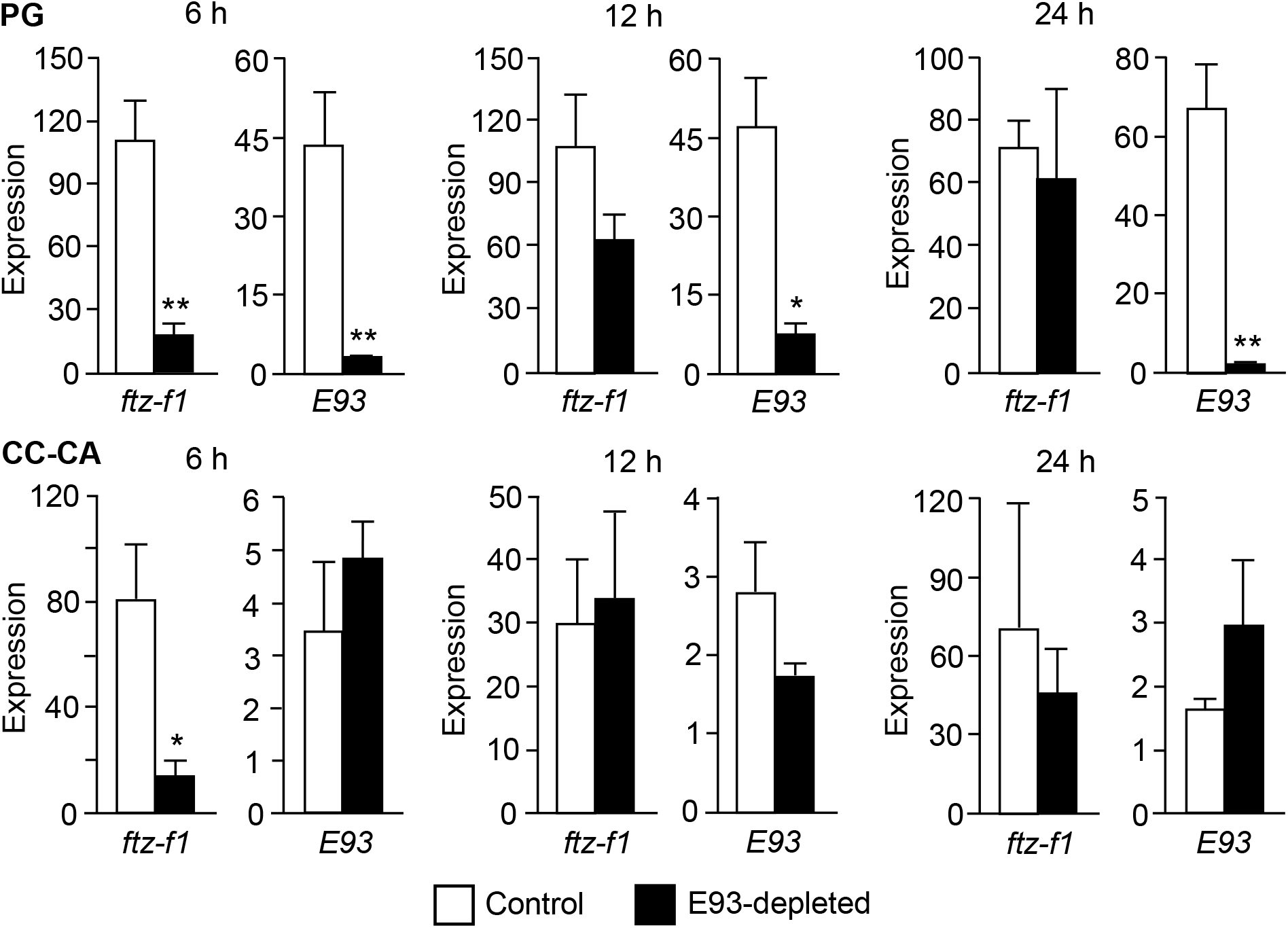
Effect of FTZ-F1 depletion on *E93* expression in *Blattella germanica*. Expression was measured in the prothoracic gland (PG), and corpora cardiaca-corpora allata (CC-CA) complex, 6, 12 and 24 h after the dsFTZ-F1 (or control) treatment in 7-day-old final instar nymphs (N6D7). (A) mRNA levels of *ftz-f1* in PG and CC-CA complex. (B) mRNA levels of *E93* in PG and CC-CA complex. The results are indicated as copies of the examined mRNA per 1000 copies of BgActin-5c mRNA, and are expressed as the mean ± SEM (n=3); the asterisks indicate statistically significant differences with respect to the controls (*p<0.05, **p<0.01) according to the student’s *t*-test.

### E93-depleted adults have transcriptionally active epidermis and wings

To characterize the type of cuticle synthesized by the E93-depleted adults, we first selected typically nymphal or typically adult cuticular proteins (NCPs and ACPs, respectively) from transcriptomic data covering the life cycle of *B. germanica*, which includes N6D6 and AdD5 (Ylla et al., 2018). As NCP genes, we selected *Bg10431, Bg10435* and *Bg15257*, which are clearly more expressed in N6 than in adults, whereas *Bg7254* and *Bg16458* were selected as ACP genes, as they are clearly more expressed in adults than in N6 (Fig. S2). The differences between N6 and adults suggested by these transcriptomic data were validated with quantitative real-time PCR (qRT-PCR) measurements (Fig. 5A). Then, measuring the NPC gene expression in AdD5 tergites showed that the levels of *Bg15257* mRNA were similar in controls and in E93-depleted insects, those of *Bg10435* tended to be higher in E93-depleted insects, whereas those of *Bg10431* were significantly higher in E93-depleted insects. The expression levels of the ACP genes were similar in E93-depleted and in controls (Fig. 5B).

**Fig. 5.**
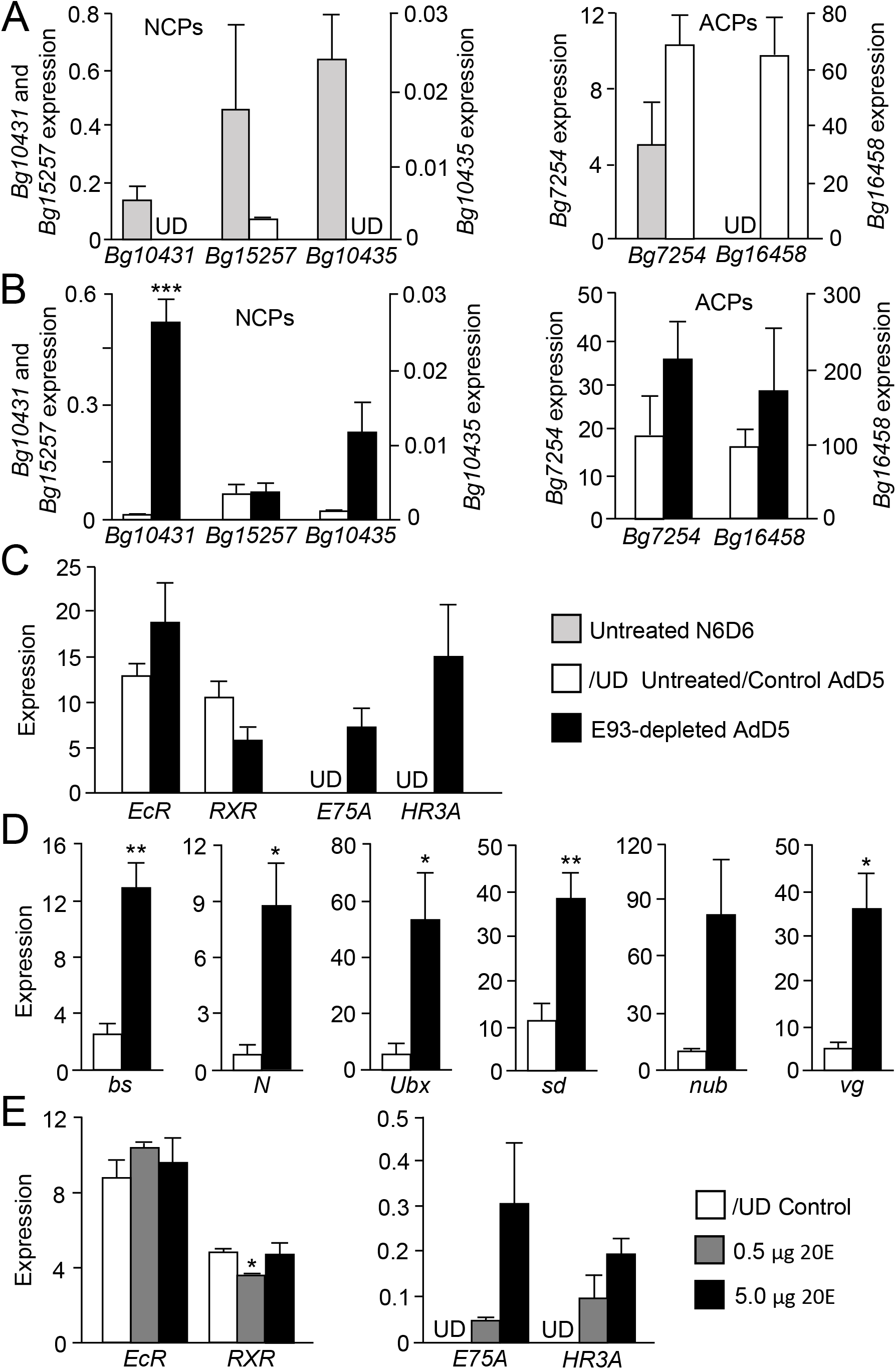
Effect of *E93* depletion and 20E treatment on gene expression in adult *Blattella germanica* tissues. (A) Expression of the nymphal cuticular protein (NCP) genes *Bg10431, Bg10435*, and *Bg15257*, and the adult cuticular protein (ACP) genes *Bg7254* and *Bg16458* in tergites in nymphs in day 6 of the sixth instar and in day 5 adults. (B) Expression of the same NCP and ACP genes in tergites of E93-depleted adults and controls in day 5 adults. (C) Expression of the ecdysteroid signaling genes: *EcR, RxR, E75A, HR3A* in tergites of E93-depleted adults and controls at day 8 of adult age. (D) Expression of wing-related genes: *blistered* (*bs*), *notch* (*N*), *nubbin* (*nub*), *scalloped* (*sd*), *ultrabithorax* (*Ubx*), and *vestigial* (*vg*) in the hindwings of E93-depleted adults and controls in day 8 adults. (E) Expression of *EcR*, *RXR, E75A* and *HR3A* in adults treated with 20E on day 8 of adult life. The results are indicated as copies of the examined mRNA per 1000 copies of BgActin-5c mRNA, and are expressed as the mean ± SEM (n=3); the asterisks indicate statistically significant differences with respect to the controls (*p<0.05, **p<0.01) according to the student’s *t*-test. In panels B and D “UD”means that mRNA levels were under the detection limit.

Next, we wondered about the differences in ecdysteroid signaling in tergites, which could explain why E93-depleted adults can molt again, whereas control adults do not. The results showed that the genes of the two components of the ecdysone receptor, *EcR* and *RXR*, are similarly expressed both in E93-depleted and control adults. In contrast, the expression of the ecdysone-dependent genes *E75A* and *HR3A* was undetectable in control adults, whereas it was clearly measurable in E93-depleted adults (Fig. 5C).

Remarkably, wings also showed signs of molting (apolysis) (Fig. 2E). We therefore looked for possible differences in the transcriptional capacity of wings from E93-depleted and control adults. To do this, we measured the expression of the following wing-related genes (Elias-Neto and Belles, 2016) in the hindwings: *blistered* (*bs*), *Notch* (*N*), *Ultrabithorax* (*Ubx*), *scalloped* (*sd*), *nubbin* (*nub*) and *vestigial* (*vg*). For all these genes, the expression was low in the controls, whereas in E93-depleted adults they were four to ten times higher, depending on the gene (Fig. 4C). Equivalent measurements carried out on the forewings gave similar results (not shown).

### Treatments with 20E do not trigger molting in control adults

Given that molting of the E93-depleted adults is associated with the preservation of the PG and active ecdysone signaling in the epidermal cells (Fig. 5C), we wondered whether exogenously administered 20E to untreated adults might trigger molting. The differences in ecdysteroid titers between nymphs and adults are of approximately one order of magnitude. The maximum levels in N6D6 are ca. 2000 pg/μl of hemolymph, whereas in 8-day-old adult females the levels are ca.125 pg/μl (Romaña et al., 1995). Thus, we injected a 0.5 μg dose of 20E in 1 μl volume into 8-day-old adult females (i.e., 2500 pg/μl of hemolymph, close to the physiological concentration in N6D6) (n=10). In another equivalent group, we injected the pharmacological dose of 5 μg of 20E in 1 μl volume (n=10). The controls received the same volume of solvent (n=10). Eight hours after the treatment, three insects of each group were used for qRT-PCR measurements, and the remaining insects were kept under observation for at least two weeks.

Expression measurements of *EcR*, *RXR, E75A* and *HR3A* in tergites showed that neither 20E treatment increased the expression of *EcR* and *RXR*. In contrast, the expression of *E75A* and *HR3A* was upregulated by 20E in a dose-dependent manner (Fig. 5E). However, even after being treated with 5 μg of 20E, the levels were at least an order of magnitude lower than in 8-day-old E93-depleted AdD8 (Fig. 5C). Importantly, none of the treated adults that we left alive showed symptoms of apolysis.

## DISCUSSION

### The expression pattern of *E93* and *ftz-f1* in the PG is tissue-specific

The expressions of *E93* and *ftz-f1* in the CC-CA, tergites, and wing pads in the last nymphal instars of *B. germanica* show a regularly fluctuating pattern, whereas in the PG there is a single expression peak of *E93* and *ftz-f1* on the last day of the final nymphal instar (Fig. 1). The characteristic *E93* and *ftz-f1* expression patters in various tissues have previously been reported in *B. germanica* (Mané-Padrós et al., 2010; Ureña et al., 2014), but our results show that there is a very peculiar stage- and tissue-specific up-regulation of these two genes in the PG, associated with the transition from the final nymphal instar to the adult. This PG-specific regulation of *E93* and *ftz-f1* is also suggested by the epistatic relationships between the two genes, as E93 does not affect the *ftz-f1* expression in the PG but represses it in the other tissues studied (Fig. 3A), whereas FTZ-F1 enhances the expression of *E93* in the PG but not in other tissues, such as the CC-CA (Fig. 4).

### E93 depletion in the PG prevents its histolysis after the imaginal molt

Given the expression patterns in the different tissues (Fig. 1), injecting dsE93 on the penultimate day of the last nymphal instar had an important effect on E93 in the PG. The result was that the insects molted to adults that were morphologically similar to the controls (Fig. 2A), but their PG did not disintegrate (Fig. 2C). The fact that the PG of the E93-depleted adults did not degenerate and were fully active was also shown by their active transcription of the steroidogenic genes (Fig. 3D). The significant expression of the ecdysone-dependent genes *E75A* and *HR3A* in the PG of E93-depleted 8-day-old adults (Fig. 3E) also supports the notion that the glands are actively producing ecdysone. It is worth noting, however, that the long-term fate of the PG in E93-depleted adults is uncertain, since this gland does not present the typical cell proliferation pattern seen in the nymphal instars (Fig. S1), as the PG undergoes cell division during the first days of each instar (Kamsoi and Belles, 2019).

Our observations revealed that E93-depletion kept the expression levels of *iap1* in the PG much higher than in the controls, when measured at the end of the final nymphal instar. Correspondingly, the expression of the effector caspase gene *casp-1* was not up-regulated in the PG of E93-depleted insects when measured on the first day of the adult stage (Fig. 3C). These E93 effects on inhibitors and promoters of PCD in the PG provide the mechanistic basis (see (Martin, 2002; Orme and Meier, 2009; Yin and Thummel, 2005; Yin et al., 2007)) for explaining why the gland did not disintegrate after the imaginal molt.

### E93 triggers PG histolysis

Previous studies reveal that FTZ-F1 depletion prevents PG histolysis in the context of *B. germanica* metamorphosis (Mané-Padrós et al., 2010). Here we show that FTZ-F1 depletion downregulates E93 in the PG (Fig. 4), suggesting that FTZ-F1 enhances the expression of E93 in this gland, which then triggers PG histolysis. This activating role of FTZ-F1 is not observed in other tissues, like the CC-CA (Fig. 4), where E93 expression is declining beyond day 6 of N6 (Fig. 1). The rapid recovery of the FTZ-F1 mRNA levels observed after the injection of dsFTZ-F1 (Fig. 4) is intriguing. Possibly, reaching high levels of FTZ-F1 expression in PG in N6D8 is crucial to undertake metamorphosis, and the gene response mechanisms to a mRNA depletion is this particular tissue and stage are especially efficient.

It is known that FTZ-F1 modulates gene expression by interacting with an FTZ-F1 response element (F1RE) located in the target gene promoter (Lavorgna et al., 1991). Interestingly, the promoter region of *B. germanica E93* contains a canonical EcR response element (Cherbas et al., 1991; Kayukawa et al., 2017), as expected, but it also contains the sequence TCAAGGCAA that might be compatible with a F1RE (De Mendonça et al., 2002; Ohno et al., 1994) (Fig. S3). This again suggests that FTZ-F1 coactivates the expression of the ecdysone-dependent gene *E93*, although this coactivating effect seems specific of the PG.

Studies on *D. melanogaster* have previously demonstrated that FTZ-F1 acts as a competence factor during prepupal development, promoting the expression of stage-specific genes like *E93* (Broadus et al., 1999; Woodard et al., 1994). Moreover, E93 has been shown to trigger PCD during metamorphosis in various tissues (salivary glands, fat body, and midgut) and species (*D. melanogaster*, and *B. mori*) (Baehrecke and Thummel, 1995; Berry and Baehrecke, 2007; Lee and Baehrecke, 2001; Lee et al., 2000; Lee et al., 2002a; Lee et al., 2002b; Liu et al., 2014; Liu et al., 2015; Wang et al., 2012; Woodard et al., 1994). Therefore, although the PCD function of E93 in the PG in the context of metamorphosis has not previously been reported, this function is not surprising given the above antecedents of E93 as a PCD effector. The role of E93 as an effector of PG histolysis after the imaginal molt supports the hypothesis that E93 may have been a crucial factor underlying the mechanisms that facilitated the evolutionary innovation of insect metamorphosis (Belles, 2019b; Belles, 2020).

### E93-depleted adults molt again

Depletion of E93 at the end of the final nymphal instar protected the PG from histolysis, but the resulting adults were morphologically normal, in general. Only14% of them presented partially unextended wings (Fig. 2A), a feature that is not uncommon even in control insects. According to the MEKRE93 pathway (Belles and Santos, 2014), the formation of morphologically normal adults is not surprising as the expression levels of *Kr-h1* (and *BR-C*) in the epidermis at the end of the final nymphal instar of E93-depleted insects are as low as in the controls (Fig. 3B). Interestingly, E93-depleted adults are able to molt again, undertaking apolysis, as shown by the double structures, notably the new mandibles, which can be observed by transparency under the old cuticle.

Even the wings of these E93-depleted adults undertake apolysis, as shown by the double veins observed, particularly in the hindwings (Fig. 2E), and they are transcriptionally active, according to the high expression levels of wing-related genes (Fig. 5C). Wing epidermal cells die and disappear after the imaginal molt, and only remain in the wing veins (Kimura et al., 2004). In E93-depleted adults we observed epidermal cells not only in the veins but also in the intervein regions. Interestingly, the newly formed veins in these adults were bigger than the old ones, thus being folded (Fig. 2E), suggesting that their epidermal cells undertook intensive cell proliferation. The high expression levels of *sd* (Fig. 5D) may have contributed to this proliferation, as this transcription factor acts together with the coactivator Yorkie, regulating Hippo pathway-responsive genes, and cell growth and proliferation (Huang et al., 2005; Zhang et al., 2008).

The cuticle of the E93-depleted adults expresses the ACP genes *Bg7254* and *Bg16458* at normally high levels, and those of the NCP gene *Bg15257* at normally low levels for an adult. However, the typically NCP genes *Bg10435* and, particularly, *Bg10431* are expressed at higher levels than normal (Fig. 5B). Thus, from the point of view of the cuticular proteins expression, the epidermis of the E93-depleted adults can be considered fundamentally adult, but with some nymphal character (Fig. 5B). In the epidermis of AdD5 insects, the expression levels of *EcR* and *RXR* were similar in E93-depleted and control insects, but the expression of *E75A* and *HR3A* was undetectable in the latter (Fig. 5C). This might suggest that adult controls cannot molt due to low levels of circulating ecdysteroids.

### Control adults do not molt even when supplied with 20E

The above conjecture led us to administer 20E to control adults, at physiological and pharmacological doses. However, even the pharmacological doses resulted only in a modest increase of *E75A* and *HR3A* expression (Fig. 5E), despite there being operative expression of *EcR* and *RXR* (Fig. 5C). Thus, the treatment did not significantly upregulate the *E75A* and *HR3A* expression levels observed in E93-depleted AdD8 insects (Fig. 5C), and did not trigger symptoms of molting. This is consistent with classically recorded experiments showing that ecdysteroids very rarely cause molting effects in adult insects, even when injected in doses as high as 3 mg/gr (Schneiderman et al., 1970). Intriguingly, however, a nymphal, active PG implanted into an adult of the Madeira cockroach *Rhyparobia* (=*Leucophaea*) *maderae* can trigger a subsequent molt, even though the insects are unable to undertake the corresponding ecdysis (Engelmann, 2002), just as in our E93-depleted *B. germanica* adults. A possible explanation for the difference between implanted PG and injected 20E is that the implanted PG releases ecdysone according to a precise pattern of increase and decrease that is crucial for triggering the appropriate cascade of gene expression (Sakurai, 2005), which would not be reproduced by an injection of 20E. Another possibility is the occurrence of a PG factor that makes the epidermal cells competent to respond to ecdysteroids and produce a new cuticle, a putative factor whose production would be interrupted with the histolysis of the PG.

A great deal of work has been done on the pupal commitment of epidermal cells in lepidopterans, including *Manduca sexta* (Hiruma et al., 1991; Riddiford, 1981) and *B. mori* (Muramatsu et al., 2008), which essentially results from the action of ecdysteroids in the absence of JH. However, the possible “adult commitment” of epidermal cells could be due to other factors, among which E93 and/or FTZ-F1 could be involved. Although RNAi targeting E93 was performed at the end of N6, when its expression levels were already declining in the epidermis and wing pads, a certain level of E93-depletion was seen in those tissues, paralleled by a dramatic upregulation of *ftz-f1* expression (Fig. 3A). It is possible therefore, that relatively high levels of E93 and/or low levels of FTZ-F1 trigger the adult commitment of epidermal cells. Whatever the case, these molting adults could represent an interesting model in which to study the mechanisms determining the adult differentiation of insect epidermal cells, and perhaps find ways to revert this process.

## MATERIALS AND METHODS

### Insects and dissections

The *B. germanica* cockroaches used in the experiments described herein were obtained from a colony fed on Panlab dog chow and water *ad libitum*, and reared in the dark at 29 ± 1^□^C and 60-70% relative humidity. Prior to injection treatments, dissections and tissue sampling (PG, CC-CA complex, abdominal tergites two to seven, and wing pads), the cockroaches were anesthetized with carbon dioxide.

### RNA extraction and retrotranscription to cDNA

RNA extractions were carried out with the Gen Elute Mammalian Total RNA kit (Sigma-Aldrich). A sample of 300 ng from each RNA extraction was used for mRNA precursors in the case of wing disc, fat body, ovary and epidermis. All the volume extracted of CC-CA complex, PG and wing was lyophilized in the freeze-dryer FISHER-ALPHA 1-2 LDplus, and then resuspended in 8 μl of milliQ H_2_O. RNA quantity and quality were estimated by spectrophotometric absorption at 260 nm in a Nanodrop Spectrophotometer ND-1000^®^ (NanoDrop Technologies). The RNA samples were then treated with DNase (Promega) and reverse transcribed with first Strand cDNA Synthesis Kit (Roche) and random hexamer primers (Roche).

### Determination of mRNA levels by quantitative real-time PCR

Measurements with qRT-PCR were carried out in an iQ5 Real-Time PCR Detection System (Bio-Lab Laboratories), using SYBR^®^Green (iTaq™ Universal SYBR^®^Green Supermix; Applied Biosystems). Reactions were carried out in triplicate, and a template-free control was included in all batches. Primers used to measure the transcripts of interest are detailed in Table S1. The efficiency of each set of primers was validated by constructing a standard curve through three serial dilutions. Levels of mRNA were calculated relative to BgActin-5c mRNA (Table S1). Results are given as copies of the examined mRNA per 1000 copies of BgActin-5c mRNA.

### RNA interference and 20E injections

The detailed procedures for RNAi assays have been described previously (Ciudad et al., 2006). The primers used to prepare the dsRNA targeting *B. germanica* E93 and FTZ-F1 are described in Table S1. The sequence corresponding to the dsRNAs (dsE93 and dsFTZ-F1) was amplified by PCR and then cloned into a pST-Blue-1 vector. A 307 bp sequence from *Autographa californica* nucleopolyhedrosis virus (Accession number K01149.1) was used as control dsRNA (dsMock). A volume of 1 μl of the dsRNA solution (3μg/μl) was injected into the abdomen of sixth instar female nymphs at the chosen ages, with a 5μl Hamilton microsyringe. Control nymphs were equivalently treated with dsMock. To study the possible molting effect of ecdysteroids in normal adults, a volume of 1 μl of a 20E solution (0.5 μg/μl or 5 μg/μl) was injected into the abdomen of 8-day-old adult females with a 5μl Hamilton microsyringe, whereas controls received 1 μl of solvent (water with 10% ethanol).

### Morphological studies of PG

Dissection of PG in female adults was carried out in Ringer’s saline. The PG was fixed in 4% paraformaldehyde for 2 h and permeabilised in PBT (0.3% Triton in PBS), then it was first incubated in 300 ng/ml phalloidin-TRITC (Sigma) for 20 min, and subsequently in 1 μg/ml DAPI (Sigma) in PBT for 5 min. After three washes with PBT, the PG was mounted in Mowiol (Calbiochem) and observed with a fluorescence microscope Carl Zeiss-AXIO IMAGER.Z1. Wings and mandibles were dissected and mounted in Mowiol. Examinations and photographs were made with a stereomicroscope Zeiss DiscoveryV8.

### Experiments to measure cell proliferation in PG

For labeling DNA synthesis and dividing cells, we followed an approach in vivo, using the commercial EdU compound “Click-it EdU-Alexa Fluor^®^594 azide” (Invitrogen, Molecular Probes). EdU was injected into the abdomen of adults at chosen ages with a 5 μl Hamilton microsyringe (1 μl of 20 mM EdU solution in DMSO). Control specimens received 1 μl of DMSO. The PG from treated specimens was dissected 1 h later, and processed for EdU visualization according to the manufacturer’s protocol.

## Acknowledgements

O.K. received a Royal Thai Government Scholarship to do a PhD thesis in X.B. laboratory, in Barcelona. We thank Jose Carlos Montañes for helping with transcriptome comparisons and searching for genes in the *B. germanica* genome, available at https://www.hgsc.bcm.edu/arthropods/german-cockroach-genomeproject, as provided by the Baylor College of Medicine Human Genome Sequencing Center. We also thank Alba Ventos-Alfonso for helping in different experiments and image treatment, and Maria-Dolors Piulachs and Jose-Luis Maestro for helpful discussions.

## Author contributions

Conceptualization: OK, X.B.; Methodology: OK, X.B.; Formal analysis: OK, X.B.; Investigation: OK, X.B.; Resources: X.B.; Writing - original draft: X.B., O.K.; Writing - review & editing: X.B.; Supervision: X.B.; Project administration: X.B.; Funding acquisition: X.B.

## Supplementary information

Supplementary information available online at

